# Adhesive Contact Between Cylindrical (Ebola) and Spherical (SARS-CoV-2) Viral Particles and a Cell Membrane

**DOI:** 10.1101/2020.06.26.173567

**Authors:** Jiajun Wang, Nicole Lapinski, X. Frank Zhang, Anand Jagota

## Abstract

A critical event during the process of cell infection by a viral particle is attachment, which is driven by adhesive interactions and resisted by bending and tension. The biophysics of this process has been studied extensively but the additional role of externally applied force or displacement has generally been neglected. In this work we study the adhesive force-displacement response of viral particles against a cell membrane. We have built two models: one in which the viral particle is cylindrical (say, representative of filamentous virus such as Ebola) and another in which it is spherical (such as SARS-CoV-2 and Zika). Our interest is in initial adhesion, in which case deformations are small and the mathematical model for the system can be simplified considerably. The parameters that characterize the process combine into two dimensionless groups that represent normalized membrane bending stiffness and tension. In the limit where bending dominates, for sufficiently large values of normalized bending stiffness, there is no adhesion between viral particles and the cell membrane without applied force. (The zero-external-force contact width and pull-off force are both zero.) For large values of normalized membrane tension, the adhesion between virus and cell membrane is weak but stable. (The contact width at zero external force has a small value.) Our results for pull-off force and zero force contact width help to quantify conditions that could aid the development of therapies based on denying the virus entry into the cell by blocking its initial adhesion.

## 1. Introduction

Virus infection is one of the major public health issues in the world. From one of the most lethal viruses, Ebola virus, to Novel Coronavirus (SARS-CoV-2) causing the present pandemic, countless people have died from virus infection and complications. Moreover, this is an ongoing concern because there will always be new viruses or mutated ones. It is therefore important to develop an understanding of how virus particles infect our body cells. One of the vital moments during infection is the internalization of the virus particle often by hijacking the normal physiological adhesive function of the receptors on the surface of the human body cell membrane.

Uptake of most virus particles into a host cell is mediated by the adhesive interaction between cell-surface and virus-surface molecules.^1–3^ Because blocking this adhesion is a prime therapeutic target, it is important to understand the mechanics of initial virus adhesion to the cell. In particular, we aim to connect adhesion properties (measured, for example, by single-molecule force spectroscopy), and the behavior of the whole virus-cell membrane interaction.

The mechanics and biophysics of this problem have been approached in a number of ways ranging from all-atom molecular dynamics simulations that contain atomistic structural detail to continuum models that contain only a few parameters that characterize virus, membrane, and adhesive properties.^4,5^ The former provides tremendous descriptive and predictive power but are, by construction, specific to a particular system. The latter allow investigation of more general principles at the cost of specificity. There is a significant literature on continuum models of adhesion between particles (biological or inorganic) and membranes. (See reference ^6^ for a review.) Shanahan developed a model for compliant particle adhesion onto a rigid surface.^7^ Long et al. modeled aspects of the fusion of synaptic vesicles with the plasma membrane, which is related to the problem of soft vesicles attaching onto a rigid surface.^8^ A similar model about kinetics of virus binding and fusion was developed by Chou.^9^ Zhang studied nanoparticle cellular endocytosis by considering parameters such as ligand density.^10^

An important subclass of models, to which our work belongs, share the feature that the total free energy is written as the sum of three contributions: from bending, tension, and adhesion. Several of the models are based on the theory that the marginal reduction of the free energy during the adhesion process balances the marginal cost of bending the membrane.^11–13^ Several studies consider the process of virus particle wrapping and engulfment by the cell membrane.^14,15^ Seifert and Lipowsky (1990) studied the adhesion of vesicles and found that there is a transition between a bound and free state based on competition between bending and adhesion energies.^11,12^ Other studies have considered the combined effects of bending, tension, and adhesion, as the particle is wrapped partially to fully.^6,13,16^

Although this problem of adhesion has been well-developed, much less attention has been paid to the contact mechanics of viruses onto a flexible membrane. This is of interest because such analyses can be used to interpret in detail atomic force microscopy (AFM) force-spectroscopy measurements of force-deflection response used to characterize the adhesion process^17^, particularly to interpret the force-spectroscopy studies on virus–host cell interactions. ^18–20^ It is also of interest because virus attachment often occurs under external load, which is accounted for in a contact mechanics model but not in most of the literature cited above. In this work, we create a continuum model for the small-deflection adhesive contact mechanics of virus particle attachment onto the host cell membrane in terms of the principal biophysical properties of the virus, membrane, and their interaction. We seek equilibrium states by minimizing the total free energy of the virus-membrane system. In particular, we seek to describe the force-deflection and contact area-deflection relationships. These results also help to retrieve conditions for lack of adhesion, pull-off force, and contact area between the virus particle and cell membrane. The common physical basis for the model is to assume that, for given applied deflection, equilibrium states are achieved by a balance of adhesion energy of cell surface receptors, which drives adhesion, against the cost of the associated deformation, represented by bending and tension energies of the cell membrane.^16,21^ We show that the solution is governed by two dimensionless materials parameters that can be identified as a normalized bending stiffness and tension. (These parameters have previously been identified.^6,12^) In particular, quantities of interest such as the equilibrium contact size (if any) or force required to remove the virus from the membrane depend (in dimensionless form) only on these two parameters. The analysis is carried out for 2D cylindrical and 3D axisymmetric virus models representing, for example, a filamentous virus like Ebola virus or a nominally spherical virus like SARS-CoV-2.

In the following sections we first outline how the problem is posed for both cylindrical and axisymmetric models. In later sections we present the results of the model. The Supporting Information contains details of derivations.

## 2. Methodology

We now describe in outline the continuum models for adhesive contact between the virus and cell-membrane, driven by adhesion and external displacement or force, and resisted by tension and elastic bending. Fig. 1 sketches the geometry of both cylindrical (i.e., 2D) and spherical (i.e., axisymmetric) models. Aspects of the 2D model have been previously studied by Mkrtchyan et al. ^22^, but without external force or displacement. For both the 2D and axisymmetric cases, we consider the case where the viral particles are stiff compared to the cell membrane to which they attach. All the parameters and non-dimensional parameters used for the models are summarized in Table 1.

**Figure 1.**
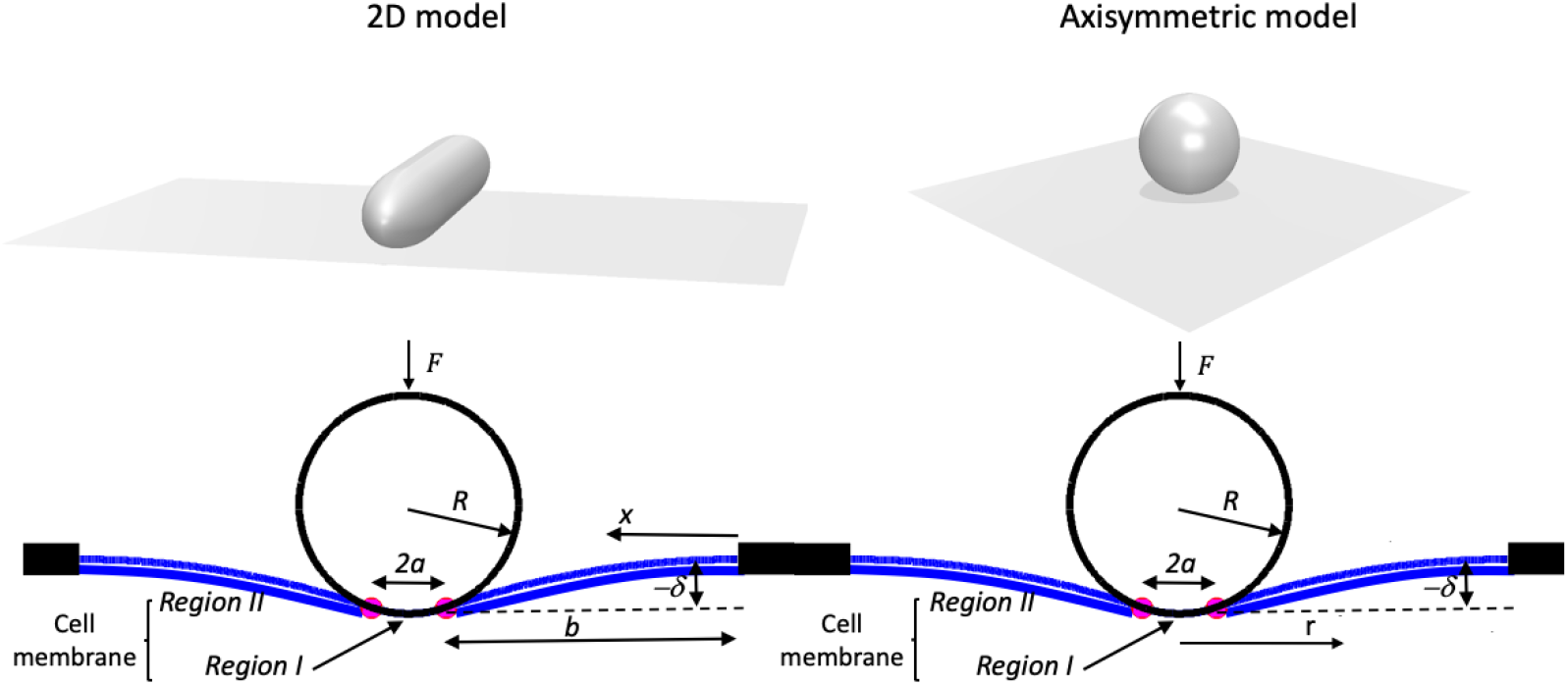
The geometry of the two models. (upper left) Sketch of a stiff cylindrical (i.e., 2D) virus particle attaching onto a flexible membrane; (upper right) Sketch of a stiff spherical virus particle attaching onto a flexible axisymmetric membrane; (lower left) mechanical model of 2D virus particle attachment driven by adhesion and external force or displacement, and resisted by membrane bending and tension; (lower right) mechanical model of spherical virus particle attachment driven by adhesion and external force or displacement, and resisted by membrane bending and tension.

**Table1.** Variables and parameters.

| Variables and Parameters | Definition | Typical Value | Normalized Variable |
| --- | --- | --- | --- |
| $\beta$ | Free energy binding per receptor. | $17 k_B T$ (TIM/PS)<sup>23</sup> | |
| $R$ | Cylindrical virus radius | $40 \text{ nm}$ (Ebola virus) | |
| $\rho$ | Density of receptors | $1000 \mu\text{m}^{-2}$ (TIM)<sup>23</sup> | |
| $\kappa$ | Cell membrane bending rigidity | $10 - 100 k_B T$ <sup>26-28</sup> | |
| $\sigma$ | Cell membrane tension | $0.1 \sim 0.3 \text{ mN/m}$ <sup>29,30</sup> | |
| $w$ | Membrane deflection | | $\bar{w} = \frac{w}{R}$ |
| $x$ | Horizontal distance from origin | | $\bar{x} = \frac{x}{R}$ |
| $a$ | Half of the contact width | | $\bar{a} = \frac{a}{R}$ |
| $l$ | Half of undeformed membrane width | | $\bar{l} = \frac{l}{R}$ |
| $b$ | $b = l - a$ | | $\bar{b} = \frac{b}{R}$ |
| $\delta$ | Virus particle indentation displacement | | $\bar{\delta} = \frac{\delta}{R}$ |
| $F$ | External force per unit length (2D model) | | $\bar{F} = \frac{F}{\rho\beta}$ |
| | External force (axisymmetric model) | | $\bar{F} = \frac{F}{\rho\beta R}$ |
| $U_{total}$ | Energy per unit length (2D model) | | $\bar{U}_{total} = \frac{U_{total}}{\rho\beta R}$ |
| | Energy (axisymmetric model) | | $\bar{U}_{total} = \frac{U_{total}}{\rho\beta R^2}$ |
| $\alpha$ | Bending stiffness normalized by adhesion | | $\alpha = \frac{\kappa}{2\rho\beta R^2}$ |
| $\gamma$ | Tension normalized by adhesion | | $\gamma = \frac{\sigma}{2\rho\beta}$ |

The cell membrane, represented in Fig. 1 by the blue line, can be split into two parts, one that contacts the virus particle and a second one that is out-of-contact. The virus particles in the 2D and axisymmetric models are assumed to be cylindrical and spherical, respectively. Because the virus is represented by a rigid cylinder, the region of the membrane in contact has a circular cross-section in 2D, ignoring end effects because the radius of the virus is quite small comparing to its length.^23^ For example, the Ebola virus has diameter of about 80 nm, whereas its length can reach 1-2 μm. In the axisymmetric model, the virus particle is assumed to be spherical and its region of contact with the membrane is a spherical cap. For example, the novel coronavirus (SARS-CoV-2) is nominally spherical with a diameter of around 120 nm. Since the diameters of both types of viruses are much smaller in size than the host cell but large compared to the thickness of the membrane (~ 4 nm), we assume that the membrane of the host cell is originally flat and deforms when in contact with the virus (blue line, Fig. 1). The width of contact section in the 2D model is 2*a* and the host cell membrane is supported some distance *l* = *a* + *b* away from the center of the virus attachment region. For nominally spherical viruses (such as HIV, Zika virus and SARS-CoV-2) the contact region is circular with radius, *a*. The membrane of the host cell is supported some radial distance *l* away from axis of symmetry. As in the 2D model, the membrane is assumed to be flat in its stress-free state; it deforms when in contact with the virus particle.

### a. Free energy calculation

For both 2D and axisymmetric models, the interaction between the viral particle and the cell membrane is driven by adhesive interactions and externally applied force or displacement. Attachment of the viral particle to the cell membrane is resisted by energy required to bend the cell membrane and by the tension it is under. We presume that the tension is set by some effect such as osmotic pressure and is not constitutively linked to the deformation, i.e., it holds a constant isotropic value. In our models, the parameters that govern the adhesive contact mechanics are (more in Table 1) bending rigid *κ*, tension *σ*, adhesion free energy per receptor, *β*, binding receptor density *ρ*, and the radius of the virus, *R*. These parameters are associated with contributions to the energy of the system: bending, tension, and adhesion (*U_bending_*, *U_tension_* and *U_adhesion_*, respectively. ^10,13,16^ There is additionally the potential of the external force or an applied external displacement. In the work presented here we carry out the contact mechanics calculation under displacement control, so it enters as a parameter. Thus, total energy is

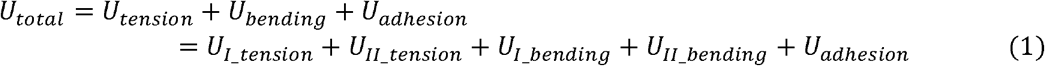

The bending and tension energies are each the sums of contributions from two regions, region I where the membrane is in contact with the virus and region II where it is free of applied lateral loads. We consider special limiting cases in which either tension or bending dominates over the other.

Calculating the energies in eq. (1) requires knowledge of the contact region and the shape of the membrane outside the contact region. This is governed by the Helfrich Hamiltonian ^13,24^ which generally results in nonlinear Euler-Lagrange governing equations.^5,8,13,25^ However, since in this work we are interested primarily in conditions close to the no-adhesion case, we can take advantage of a major simplification in the shape equation for small deformations. We can additionally neglect the coefficient of Gaussian curvature in the Helfrich Hamiltonian because the processes we study involve no change in topology. With these two simplifications, the governing equation simplifies to a linear differential equation:^13^

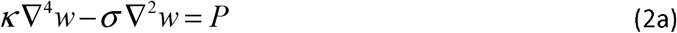

Typical values are tension *σ*~μN/m, bending rigidity *κ*~10–100 *k_B_T* ^5,10,24^, where *k_B_* is Boltzmann’s constant. In the following we will use specific forms of (2a) for plane and axisymmetric problems. The ratio 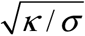 represents a length scale. If the size of the virus is large compared to this length scale, then tension dominates. Conversely, if the virus is small compared to this length, then bending does. For the viral particles we wish to consider, using typical values given in Table.1, we estimate this ratio to be about 50 nm, which means that both tension and bending energies can be important. A normalized version of equation (2a) is

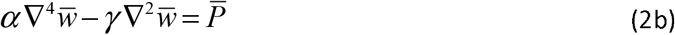

where, 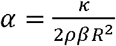 is a normalized bending stiffness, 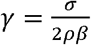 is normalized tension, *ρβ* as the product of number density of surface ligand-receptor pairs and the adhesion energy of one such pair, is the specific (per unit area) adhesion energy of the interface. Thus, in normalized form this problem is governed by two dimensionless materials parameters, *α* and *γ*. Versions of these two parameters are well-known in the literature.^6,12,13^

### b. 2D and Axisymmetric Governing Equations

In the 2D model, the membrane in region I conforms to the cylindrical virus particle so that the normalized deflection *w*(*x*) in region I is governed by the circular shape of the virus cross-section. The deflection *w*(*x*) in region II governed by the 1D version of eq. (2a):^16^

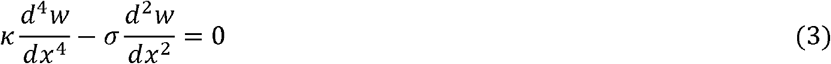

Normalizing equation (3) as defined in Table 1, we get

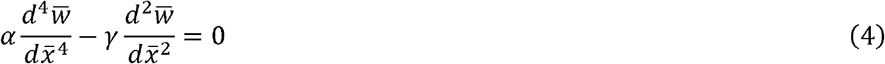

To better understand the behavior, we focus most of our attention on the two limits in which either tension or bending dominate. (We can ignore bending if 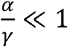; we can ignore tension if 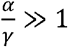.)

In the axisymmetric model, the deflection *w*(*x*) in region I is constrained by contact of the membrane to the surface of the viral particle. The problem is axisymmetric and so deflection *w*(*x*) is a function or radial position only. In region II it is governed by the differential equation:^16^

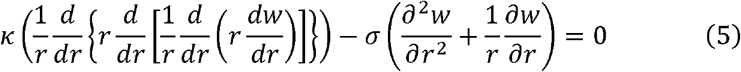

where *r* is the radial distance from the lower pole. Table 1 describes the variables and parameters, in dimensionless form.

The normalized form governing equation for the axisymmetric model is

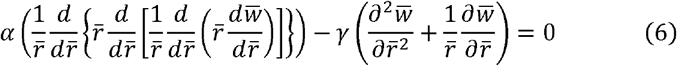

Again, we focus on the two limits in which either tension or bending dominate. Also, for the majority of the discussion in the remainder of this manuscript when we refer, for example, to tension, we mean normalized tension, since the discussion is almost exclusively of solutions to the normalized dimensionless governing equations and parameters.

### c. Solution Procedure

In normalized variables, the state of the system is described by five dimensionless variables (under displacement control): 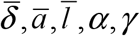, i.e., the normalized displacement of the virus, contact width/radius, size of the membrane patch, normalized bending stiffness, and tension. We find that the effect of 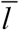 is weak and we hold it fixed in this work at 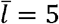, thus reducing to four the number of dimensionless variables. In region I, the deflection is specified by the fixed shape of the virus, and 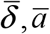. In region II, the deflection of the membrane is given by solving equation (4) or (6) subject to clamped boundary conditions where the membrane is held fixed and continuity and smoothness of deflection at the intersection between the two regions. For example, in the axisymmetric spherical virus case where bending dominates, the deflection in region II is given by the solution of equation (6) and is

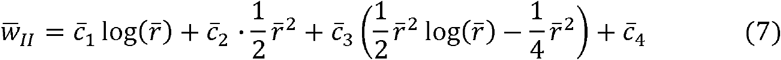

where the constants *c_1_*, *c_2_*, *c_3_*, *c_4_* are obtained by the boundary and continuity conditions (see SI for details). With the solution for deflection known, the various contributions to energy can be computed in normalized form. Again, for the same case,

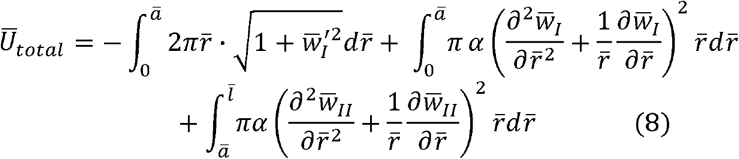

where the first term represents adhesion energy (negative in value), the second term the bending energy inside the contact region, and the third term the bending energy in region II. In general, the total energy associated with a particular configuration of the system can be written as

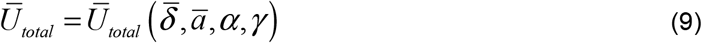

We now impose two equilibrium conditions. The first enforces the condition that for other fixed parameters, the contact radius adjusts to satisfy the condition that

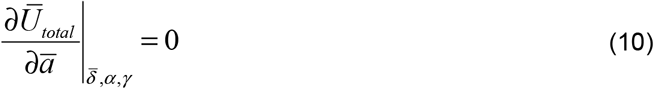

When this condition can be satisfied, it provides a solution 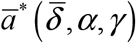, eliminating the explicit dependence of total energy on contact width so that 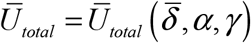. (This condition can sometimes be satisfied exactly; on other occasions it is computed numerically.) Next we apply the condition that the force is conjugate to the applied deflection so that

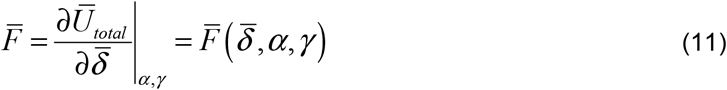

In this way, for all the different conditions studied, we obtain force-deflection 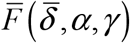 and contact width-deflection 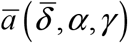 relations. A special point in the force-deflection trace is the maximum or pull-off force, 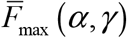. Similarly, an important point in the contact width-deflection trace is contact width corresponding to zero force, 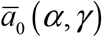. Both these quantities depend only on the two materials parameters. For the sake of brevity, those results are presented in SI.

The discussion just presented to obtain the solution in each of the cases studied is captured by the algorithm described below:

#### ALGORITHM

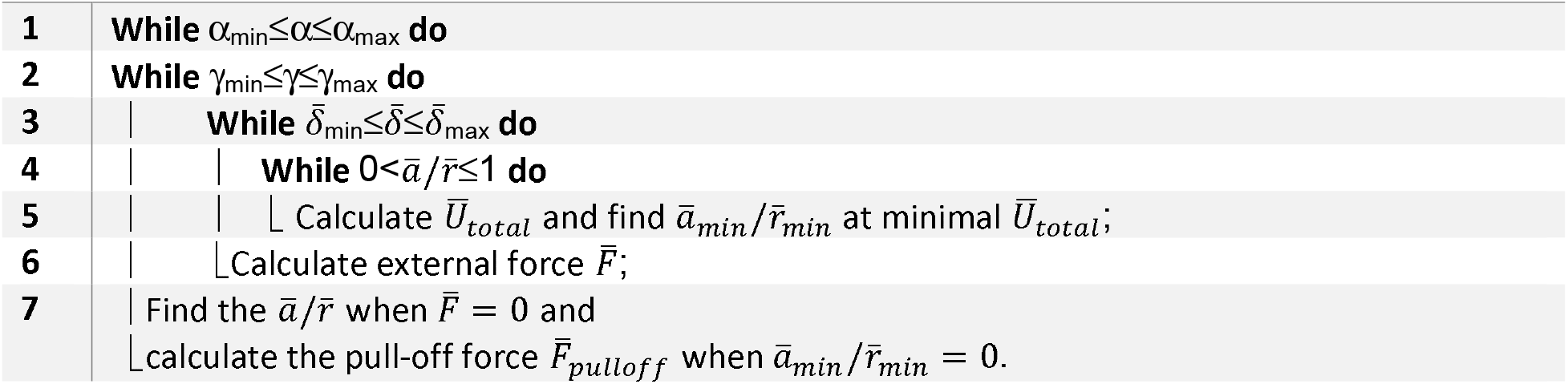

## 3. Results and Discussion

We first present results for the cylindrical virus (Figure 2). Fig. 2(a,b) show results in the bending dominated limit; Figs 2(c,d) for the tension-dominated limit. The tension-dominated limit can also be solved exactly without the small-deflection assumption. For consistency, in the main text we present results for small deflections only. The exact result for this specific case is provided in Supporting Information S1.1.

**Figure 2.**
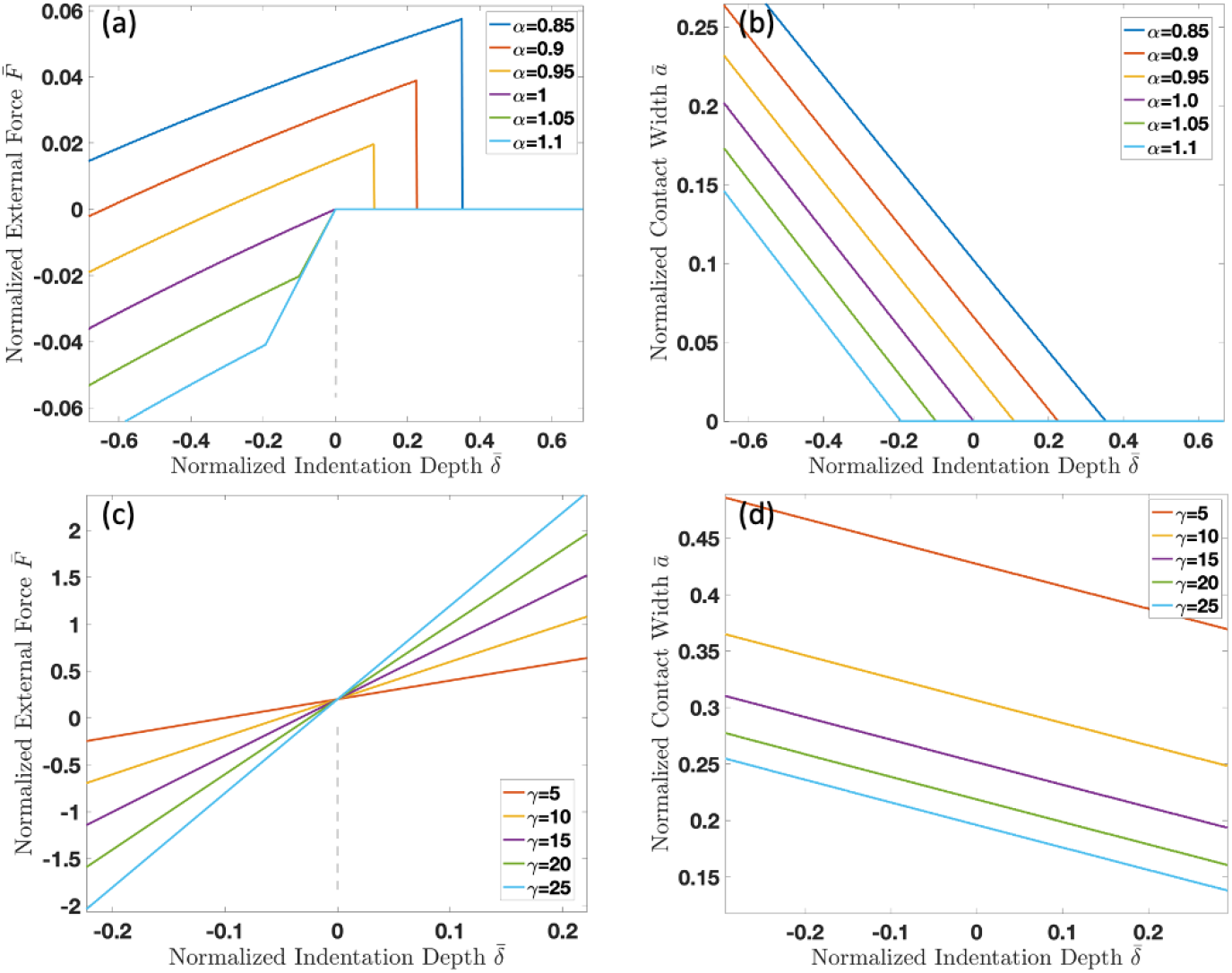
Cylindrical virus in contact with a membrane. (a) Force and (b) contact width as a function of indentation depth in the bending-dominated limit for various values of *α*. (c) Force and (d) contact width as a function of indentation depth in the tension-dominated limit for various values of *γ*.

Positive values of force represent tension. In the bending dominated limit, the force-deflection relation is approximately (but not exactly) linear. Its most interesting feature is a maximum in the force, (for *α*<1) which corresponds (Fig. 2b) to a reduction in contact width down to zero. With increasing bending stiffness *α* the peak force reduces. At the critical value of *α=1*, indicated by the dashed vertical lines, the pull-off force reduces down to zero. That is, there is no adhesion between the virus and cell membrane in the absence of external force. This result is consistent with conclusions obtained by models based only on the balance of bending and adhesion.^23^ For higher values of bending stiffness, the contact reduces to zero while still under compressive load. Subsequently, the force-deflection behavior follows that of a cylinder in line contact with the membrane. Of course, in reality, true line contact is not possible due to the finite thickness of the membrane.

Fig 2(c) shows the force-displacement results in the tension-dominated limit for a range of *γ*. The force-deflection response is nearly linear again, with slope that increases with increasing tension. Interestingly, the force for all values of *γ* is identical at a positive (tensile) value for zero indentation. As a consequence, indentation depth at zero force is negative and increases with decreasing *γ*. The contact radius as a function of indentation depth is shown in Fig. 2(d). Unlike the bending dominated case, even for large *γ*, the contact width is quite substantial for considerable tensile extension. This is unlike the bending-dominated case in which there is no force-free adhesion if bending stiffness exceeds a critical value. This means that for the tension-dominated limit of the 2D model, there isn’t a critical value at which adhesion is blocked – see also Figure S1 in Supporting Information.

Figure 3 shows results for the spherical virus adhesion. Figs 3(a,b) shows the force-deflection and contact radius-deflection results in the bending-dominated limit. Like in the 2D model, for *α>1*, the contact reduces to zero while deflection is still negative. For this range of bending stiffness, contact radius at zero force is zero as is the pull-off force (See SI). Fig. 3 (c) shows force-deflection results for the tension-dominated limit. The results here are strikingly different from the 2D model in that the force jumps to zero, corresponding to a jump to zero in the normalized contact radius, Fig 3(d). The pull-off force reduces with increasing tension, but only mildly so.

**Figure 3.**
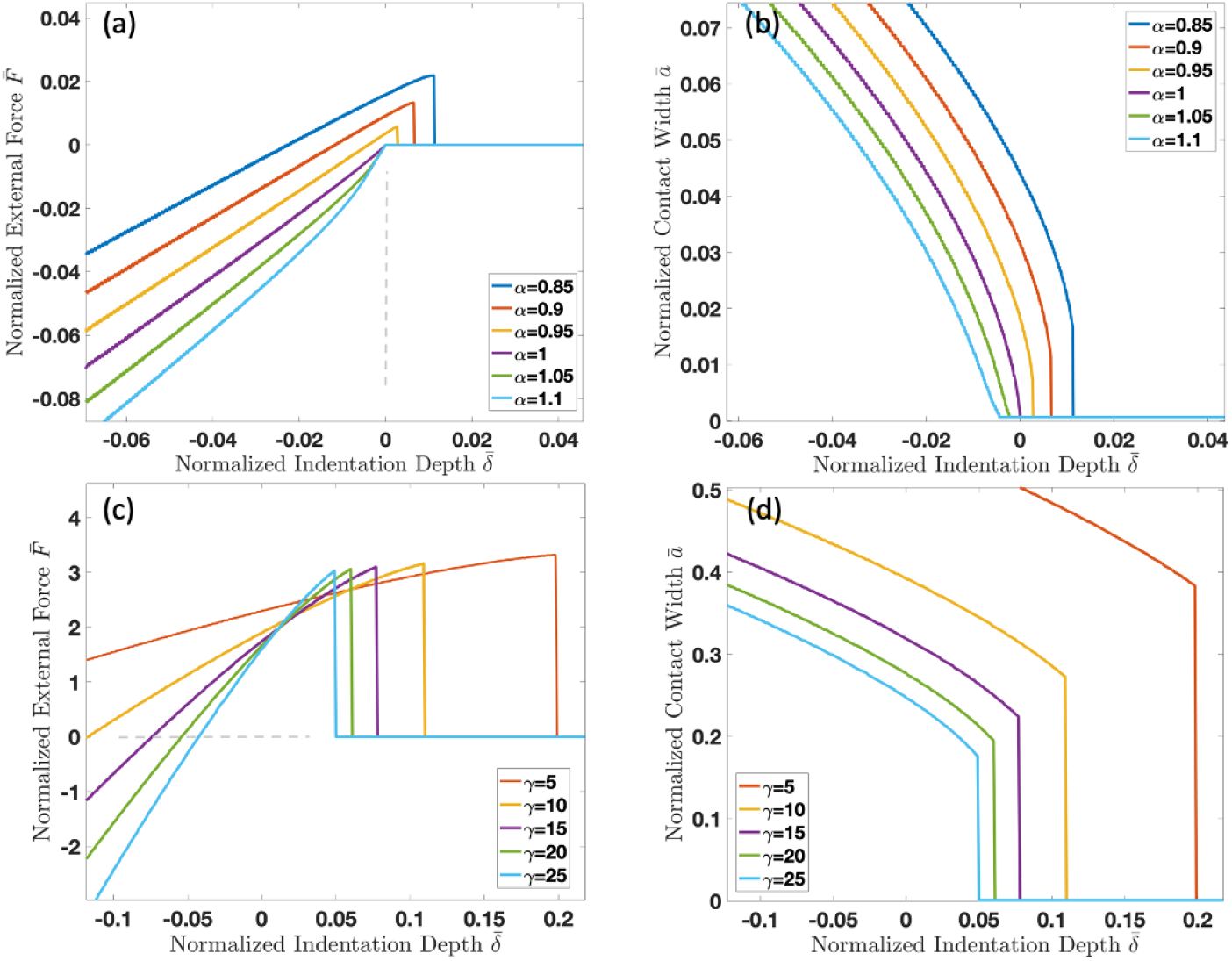
(a) Force-indentation of the spherical virus under bending dominated limit. (b) Contact radius as a function of indentation in the bending dominated limit. (c) Force-deflection in the tension dominated limit, (d) contact width in the tension dominated limit.

## 4. Conclusions

We developed and studied adhesive contact between a stiff viral particle and a cell-membrane. Our model is based on a continuum description: minimization of energy that is composed of contributions from bending, tension, adhesion, and external force or deflection. We considered cylindrical (filamentous) and spherical viruses, representative of two typical viral shapes such as Ebola and SARS-CoV-2. The principal result of the analyses is normalized force-deflection and contact width/radius versus deflection as a function of two dimensionless materials parameters, *α* and *γ*, denoting normalized bending stiffness and tension, respectively. We paid attention to limits in which bending, or tension dominate the other.

In both the bending-dominated cases, a striking result is that for sufficiently stiff membranes there is zero pull-off force and contact radius at zero force also vanishes. For larger stiffness, contact vanishes even under compressive indentation. The tension-dominated case is qualitatively different for the 2D and axisymmetric shapes. In the tension-dominated limit of the cylindrical model, the contact width 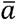 decreases very slowly with increasing deflection. In the axisymmetric geometry, in contrast, there is a well-defined pull-off force that corresponds to contact shrinking to zero. We believe that the force-displacement results can be related directly to the atomic force microscopy (AFM) force-displacement measurements. ^18 19,20^

Our continuum model is a simple representation, attempting to capture many important details in only a few physical parameters. In vivid biological context, things will be more complicated. However, our model is suitable for application to interpretation of nanoindentation and force spectroscopy experiments. It also advances understanding of the biomechanics of filamentous and spherical virus adhesive contact mechanics to cell membranes.

## Acknowledgements

This work was supported by the NIH grant 1 R15 AI133634-01A1 and by NSF grant 1804117.

