## Supplementary Information for "Adhesive Contact Between Cylindrical (Ebola) and Spherical (SARS-CoV-2) Viral Particles and a Cell Membrane"

**S1. Details of the Cylindrical Model (model for Ebola Virus)**

In this section we provide details for the 2D/Cylindrical virus models.

**S1.1 Exact solution in the tension-dominated limit**

In nearly all the cases studied in this work, we have made use of the small deflection assumption that results in linear governing differential equations for the membrane shape (eqs 2-6 in the main text). However, we begin by solving a special simple case in which we can find the exact solution without this restriction.

Figure S1 show a cylindrical virus in adhesive contact on an initially flat membrane. Only the symmetric right half of the membrane is drawn.

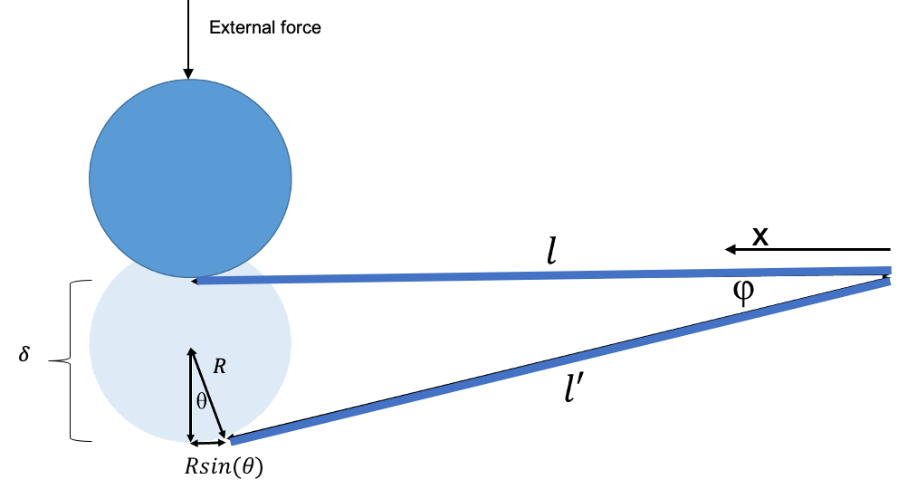

Figure S1. Geometry of the cylindrical model of a virus adhering to a membrane under constant tension.

The vertical displacement of the virus particle with respect to the flat part of the membrane is *δ*. The length of the membrane in the model is $l$ when there is no adhesion ($x=0$ at right end and $x=l$ at left end). The membrane comprises two parts: regions I and II. Region I is where the membrane is in contact with the virus particle. The rest of the membrane comprises region II. With increasing membrane deflection its length increases. For example, the length of region II in Fig. S1 increases to $l'$. Let the angle between segments labelled $l$ and $l'$ be ϕ. The radius of the virus is *R* and the angle corresponding to the arc of region I is *θ*. From simple geometry, $\cos\left( \emptyset\right)=\frac{l-Rsin\theta}{l'}$, or $l^{'}=\frac{l-Rsin(\theta)}{\cos\left( \emptyset\right)}$. Under displacement control, the total energy of the system can be calculated by summing the adhesion ($U_{adhesion}$) and tension energies ($U_{tension}$):

$$U_{total}=U_{tension}+U_{adhesion}=T\left( \frac{l-Rsin\left( \theta\right)}{\cos\left( \emptyset\right)}+R\theta\right)-\rho\beta R\theta(S1)$$

The tension energy, calculated as tension $T$ times the membrane length after deflection, is positive while the adhesion energy, calculated as the product of length of region I, energy per bond, and bond density, is negative. By geometry, can write $\emptyset$ in terms of $\theta$, thus eliminating the former:

$$\delta-(R-Rcos\left( \theta\right))=\left( \frac{l-R\sin\left( \theta\right)}{\cos\left( \emptyset\right)} \right)\sin\left( \emptyset\right) (S2)$$

so that

$$\emptyset\left( \theta\right)=\tan^{-1} \left( \frac{\delta-R+\mathrm{Rcos} \left( \theta\right)}{l-\mathrm{Rsin} \left( \theta\right)} \right) (S3)$$

So, the energy equation is

$$U_{total}=U_{tension}+U_{adhesion}=T\left( \frac{l-Rsin\left( \theta\right)}{\cos\left( \tan^{-1} \left( \frac{\delta-R+\mathrm{Rcos} \left( \theta\right)}{l-\mathrm{Rsin} \left( \theta\right)} \right) \right)}+R\theta\right)-\rho\beta R\theta(S4)$$

In dimensionless form, this is:

$$\bar{U}_{total}=\bar{U}_{tension}+\bar{U}_{adhesion}=\gamma\left( \left( \frac{\bar{l}-\sin\left( \bar{\theta} \right)}{\cos\left( \tan^{-1} \left( \frac{\bar{\delta}-1+\cos\left( \bar{\theta} \right)}{l-\sin\left( \bar{\theta} \right)} \right) \right)} \right)+\bar{\theta} \right)-\bar{\theta} (S5)$$

and the external force on the membrane is

$$\bar{F}=2\gamma sin(\emptyset(\delta)) (S6)$$

**S1.2 Small Deflection Model in the Tension-Dominated Limit**

The normalized governing equation for deflection in region II is (See Fig. 1a in main text)

$$\alpha\frac{d^{4}\bar{w}}{d\bar{x}^{4}}-\gamma\frac{d^{2}\bar{w}}{d\bar{x}^{2}}=0 (S7)$$

In the tension-dominated limit, $\frac{\alpha}{\gamma}\ll1$, and this simplifies to

$$\gamma\frac{d^{2}\bar{w}}{d\bar{x}^{2}}=0 \left( S8 \right)$$

Region I (Virus-Membrane Adhesion)

The normalized membrane deflection and its derivative in region I are

$$\bar{w}_{I}\left( \bar{x} \right)=\bar{\delta}+\frac{\left( \bar{l}-\bar{x} \right)^{2}}{2} \left( S9 \right)$$

$${\bar{w}_{I}}^{'}\left( \bar{x} \right)=-\left( \bar{l}-\bar{x} \right) \left( S10 \right)$$

Their values at the contact edge are

$$\bar{w}_{I}\left( \bar{b} \right)=\bar{\delta}+\frac{\left( \bar{l}-\bar{b} \right)^{2}}{2} \left( S11 \right)$$

$${\bar{w}_{I}}^{'}\left( \bar{b} \right)=-\left( \bar{l}-\bar{b} \right) \left( S12 \right)$$

Region II (No Virus-Membrane Adhesion)

Integrating (S8) gives:

$$\bar{w}_{II}=\bar{c}_{1}\bar{x}+\bar{c}_{2} \left( S13 \right)$$

The deflections must satisfy the boundary and matching conditions:

$$\bar{w}_{II}\left( 0 \right)=0 (S14)$$

$$\bar{w}_{II}\left( \bar{b} \right)=\bar{w}_{I}\left( \bar{b} \right)=\bar{\delta}+\frac{\left( \bar{l}-\bar{b} \right)^{2}}{2}=\bar{c}_{1}\bar{b} (S15)$$

which yield

$$\bar{c}_{1}=\frac{\bar{\delta}}{\bar{b}}+\frac{1}{2\bar{b}}\left( \bar{l}-\bar{b} \right)^{2} (S16)$$

$$\bar{c}_{2}=0 (S17)$$

Using the fact that the change in length of a membrane under tension is ~ $\left( w' \right)^{2}/2$:

$$U_{total}=U_{tension}+U_{bending}=\int_{0}^{b} \frac{1}{2}\sigma\left( w_{II}^{'} \right)^{2}dx+\int_{b}^{l} \frac{1}{2}\sigma\left( w_{I}^{'} \right)^{2}dx-\rho\beta\left( l-b \right) (S18)$$

The normalized energy is:

$$\bar{U}_{total}=\bar{U}_{tension}+\bar{U}_{adhesion}=\int_{0}^{\bar{b}} \gamma\left( \bar{w}_{II}^{'} \right)^{2}d\bar{x}+\int_{\bar{b}}^{\bar{l}} {\gamma\left( \bar{w}_{I}^{'} \right)}^{2}d\bar{x}-\left( \bar{l}-\bar{b} \right)=\gamma{{(\bar{c}}_{1}}^{2}\bar{b}+\frac{{(\bar{l}-\bar{b})}^{3}}{3})-\left( \bar{l}-\bar{b} \right) (S19)$$

Location of the contact edge, $\bar{b}$, is given by minimizing $\bar{U}_{total}$ with respect to $\bar{b}$. The force is then given by

$$\bar{F}=\gamma\frac{d\bar{w}}{d\bar{x}}=\gamma\bar{c_{1}} (S20)$$

The results for force and contact width as a function of applied deflection are shown in Figure 2(c) of the main text. Figure S1 shows contact width at zero force
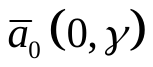
. Note that even for very large tension, the contact width decreases quite slowly. In particular, there is no specific pull-off force. Contrast this with the case presented later where bending dominates.

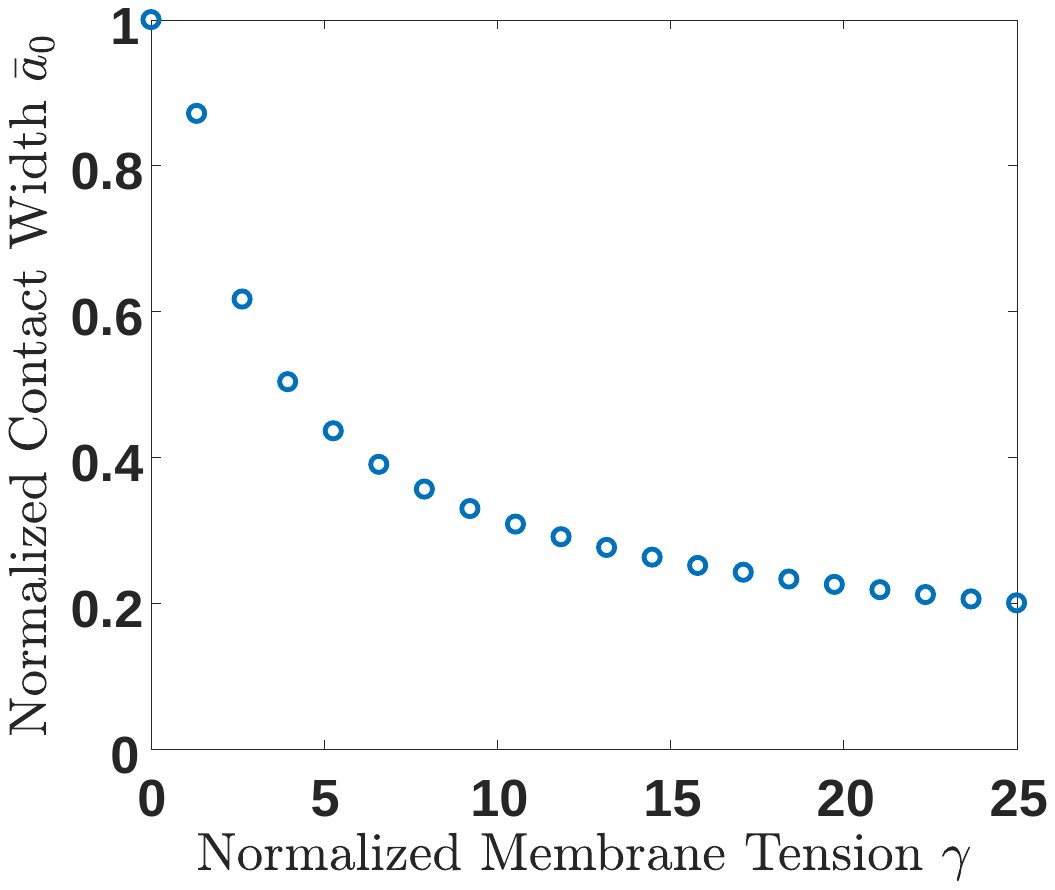

Figure S1. Normalized contact width at zero force as a function of normalized membrane tension. Contact width decreases slowly with increasing tension. In particular, unlike the bending dominated case, there is no critical value of tension at which adhesion is blocked.

**S1.2 Bending-Dominated Limit**

In this case, $\frac{\alpha}{\gamma}\gg1$ and so the normalized governing equation for membrane shape reduces to

$$\alpha\frac{d^{4}\bar{w}}{d\bar{x}^{4}}=0 (S21)$$

The normalized membrane deflection and its derivative in region I are:

$$\bar{w}_{I}\left( \bar{x} \right)=\bar{\delta}+\frac{\left( \bar{l}-\bar{x} \right)^{2}}{2} (S22)$$

$${\bar{w}_{I}}^{'}\left( \bar{x} \right)=-(\bar{l}-\bar{x}) (S23)$$

Their value at the contact edge is:

$$\bar{w}_{I}\left( \bar{b} \right)=\bar{\delta}+\frac{\left( \bar{l}-\bar{b} \right)^{2}}{2} (S24)$$

$${\bar{w}_{I}}^{'}\left( b \right)=-(\bar{l}-\bar{b}) (S25)$$

Integrating the governing equation gives:

$$\bar{w}_{II}=\bar{c}_{1}+\bar{c}_{2}\bar{x}+\bar{c}_{3}\bar{x}^{2}+\bar{c}_{4}\bar{x}^{3} (S26)$$

The matching conditions at $\bar{x}=b$ and boundary conditions at $\bar{x}=0$:

$$\bar{w}_{II}\left( \bar{b} \right)=\bar{w}_{I}(b)=\bar{\delta}+\frac{\left( \bar{l}-\bar{b} \right)^{2}}{2} (S27)$$

$${\bar{w}_{II}}^{'}\left( b \right)={\bar{w}_{I}}^{'}\left( b \right)=-(\bar{l}-\bar{b}) (S28)$$

$$\bar{w}_{II}(0)=0 (S29)$$

$${\bar{w}_{II}}^{'}(0)=0 (S30)$$

Solving S27-S29 for the unknown constants we get:

$\bar{c}_{1}=0$, $\bar{c}_{2}=0$, $\bar{c}_{3}=\frac{\bar{b}^{2}+6\bar{\delta}-4\bar{bl}+3\bar{l^{2}}}{2\bar{b^{2}}}$, $\bar{c}_{4}=\frac{-2\bar{\delta}+(\bar{b}-\bar{l})\bar{l}}{\bar{b^{3}}}$	(S30a)

The normalized total energy can be expressed as:

$$\bar{U}_{total}=\bar{U}_{elastic}+\bar{U}_{adhesion}=\bar{U}_{elasticI}+\bar{U}_{elasticII}+\bar{U}_{adhesion}=\int_{\bar{b}}^{\bar{l}} \alpha{({\bar{w}^{''}}_{I})}^{2}dx+\int_{0}^{\bar{b}} \alpha{({\bar{w}^{''}}_{II})}^{2}dx-(\bar{l}-\bar{b}) (S31)$$

Integrating this equation yields:

$$\bar{U}_{Total}=\alpha\left( \bar{l}-\bar{b} \right)+\alpha\left( 4\bar{c}_{3}^{2}\bar{b}+12\bar{c}_{4}^{2}\bar{b}^{3}+12\bar{c}_{3}\bar{c}_{4}\bar{b}^{2} \right)-(\bar{l}-\bar{b}) (S32)$$

Again, we find the equilibrium point where $\frac{d\bar{U_{T}}}{d\bar{b}}=-\alpha+\alpha\left( 4\bar{c}_{3}^{2}+36\bar{c}_{4}^{2}\bar{b}^{2}+24\bar{c}_{3}\bar{c}_{4}\bar{b} \right)+\bar{b}=0$, then loop over δ. To find the shear force

$$\bar{F}=-2\alpha\frac{d^{3}\bar{w}}{d\bar{x}^{3}}=-12\alpha\bar{c}_{4} (S33)$$

The results for force and contact width as a function of applied deflection are shown in Fig. 2(a,b) of the main text. As noted there, at a critical value of *α=1*, there is no pull-off force. Figure S2(a) below shows the contact width at zero applied force decreases with increasing bending stiffness and, goes to zero at the critical value. The pull-off force, Fig. S2(b) shows a similar and consistent trend.

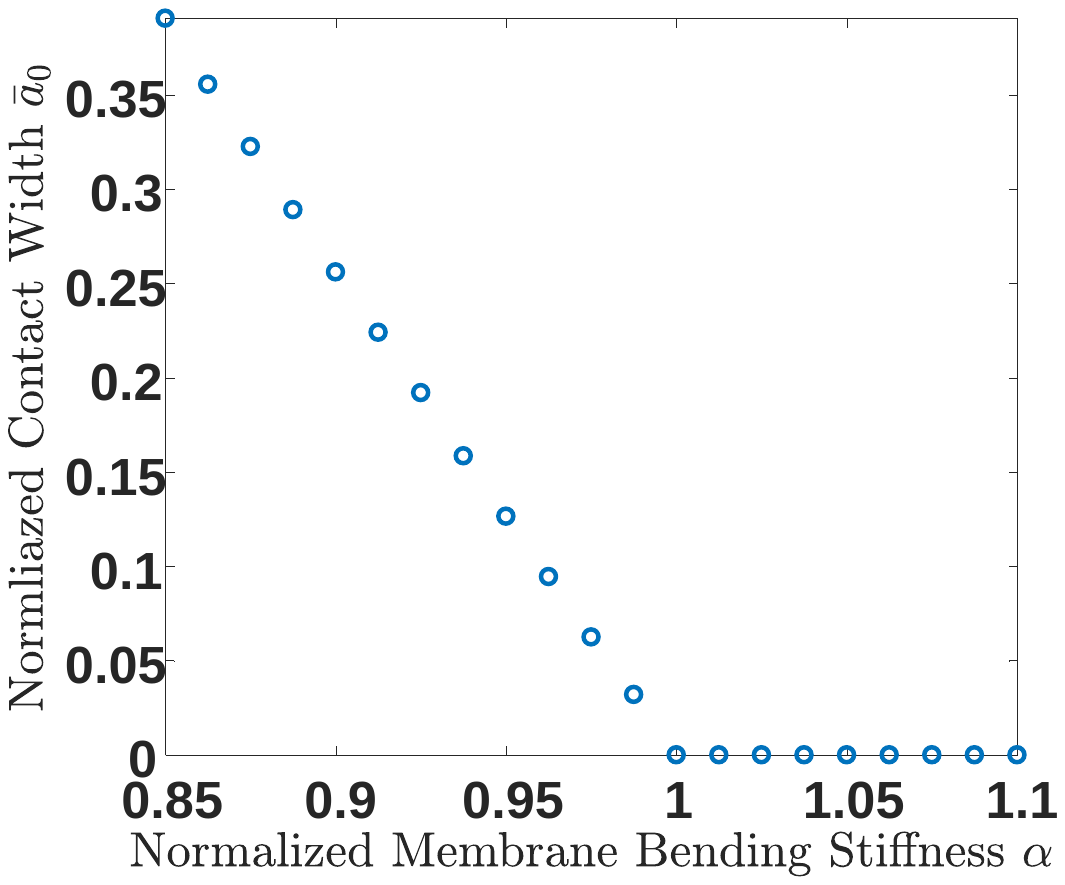

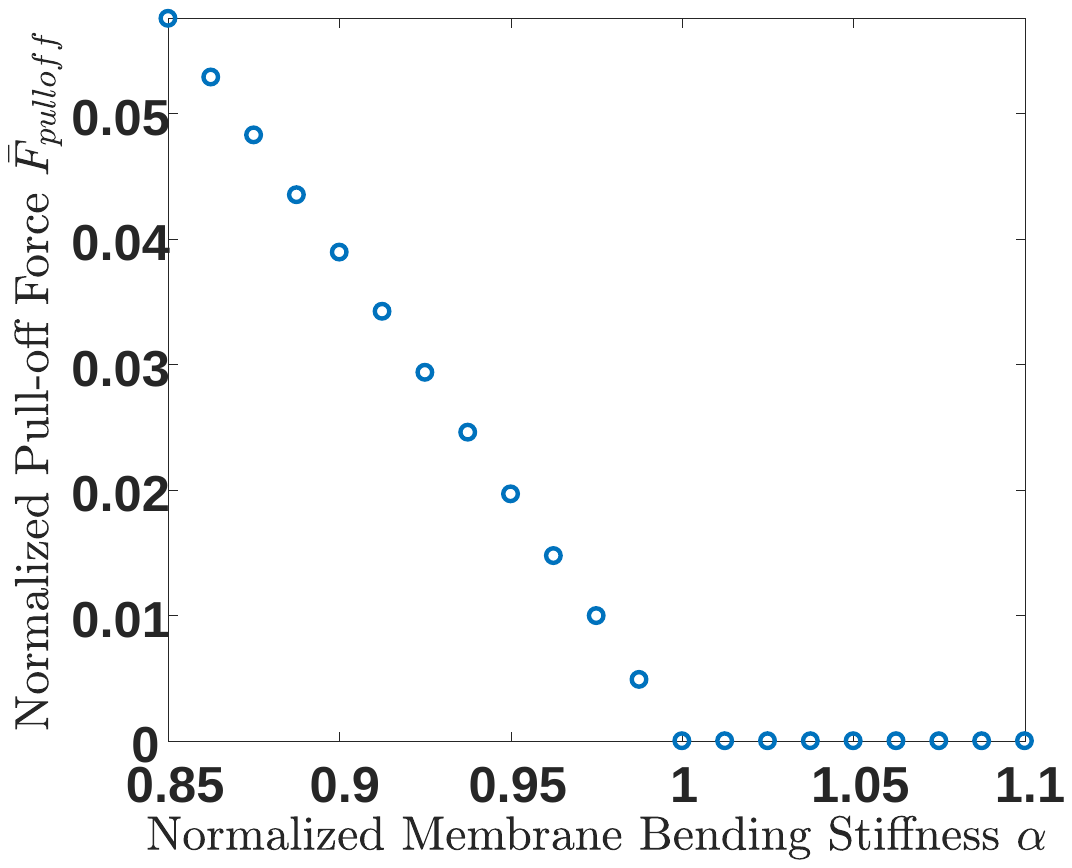

(a)	(b)

Figure S2 Plots of (a) contact width at zero force, and (b) pull-off force, both as a function of normalized bending stiffness.

**S1.3 Bending and Tension**

When neither bending nor tension can be neglected, the governing equation remains eq. (4) of the main text. The deflection of region I remains the same:

$$\bar{w}_{I}\left( \bar{x} \right)=\bar{\delta}+\frac{\left( \bar{l}-\bar{x} \right)^{2}}{2} \left( S34 \right)$$

whereas the solution of the governing equation is now

$$\bar{w}_{II}\left( \bar{x} \right)= \bar{c}_{1}+\bar{c}_{2}\bar{x}+\bar{c}_{3}\exp\left( -\sqrt{\frac{\gamma}{\alpha}}\bar{x} \right)+\bar{c}_{4}\exp\left( \sqrt{\frac{\gamma}{\alpha}}\bar{x} \right) \left( S35 \right)$$

Then applying the four boundary and matching conditions (S27-S30), we can determine the four parameters $\bar{c}_{1}, \bar{c}_{2}, \bar{c}_{3}, \bar{c}_{4}$. Then the total energy on the membrane is calculated by

$$\bar{U}_{total}=\bar{U}_{tension}+\bar{U}_{elastic}+\bar{U}_{adhesion}=\int_{\bar{b}}^{\bar{l}} \gamma\left( \bar{w}_{I}^{'} \right)^{2}dx+\int_{0}^{\bar{b}} \gamma\left( \bar{w}_{II}^{'} \right)^{2}d\bar{x}+\int_{\bar{b}}^{\bar{l}} \alpha\left( \bar{w}_{I}^{''} \right)^{2}d\bar{x}+\int_{0}^{\bar{b}} \alpha\left( w_{II}^{''} \right)^{2}dx-\left( \bar{l}-\bar{b} \right) (S36)$$

And the normalized external force is

$$\bar{F}=-2\alpha\frac{d^{3}\bar{w}}{d\bar{x}^{3}}+\gamma\frac{d\bar{w}}{d\bar{x}} (S37)$$

**S2. Details of axisymmetric model (model for SARS-CoV-2)**

**S2.1 Tension-dominated limit**

The governing equation is obtained from equation (6) of the main text by setting *α=0*:

$$\frac{\partial^{2}\bar{w}}{\partial\bar{r}^{2}}+\frac{1}{\bar{r}}\frac{\partial\bar{w}}{\partial\bar{r}}=0 (S38)$$

The solution in Region I is unchanged:

$$\bar{w}_{I}\left( \bar{r} \right)=\bar{\delta}+\frac{\bar{r}^{2}}{2} (S39)$$

$$\bar{w}_{I}'\left( \bar{r} \right)=\bar{r} (S40)$$

In region II, we obtain the solution by integrating (S38) twice:

$$\bar{w}\left( \bar{r} \right)=\bar{C}_{1}\ln\left( \bar{r} \right)+\bar{C}_{2} (S41)$$

Applying the boundary and matching conditions:

$$\bar{w_{I}}\left( \bar{a} \right)=\bar{\delta}+\frac{\bar{a}^{2}}{2} (S42)$$

$$\bar{w_{I}}\left( \bar{l} \right)=0 (S43)$$

yields:

$$\bar{w}\left( \bar{r} \right)=\bar{C}_{1}\ln\left( \bar{r} \right)+\bar{C}_{2}=\frac{\bar{\delta}+\frac{\bar{a}^{2}}{2}}{\ln\left( \bar{a} \right)-\ln\left( \bar{l} \right)}\ln\left( \bar{r} \right)+\frac{\bar{\delta}+\frac{\bar{a}^{2}}{2}}{\ln\left( \bar{l} \right)-\ln\left( \bar{a} \right)}ln(\bar{l}) (S44)$$

where

$$\bar{C}_{1}=\frac{\bar{\delta}+\frac{\bar{a}^{2}}{2}}{\ln\left( \bar{a} \right)-\ln\left( \bar{l} \right)}, \bar{C}_{2}=\frac{\bar{\delta}+\frac{\bar{a}^{2}}{2}}{\ln\left( \bar{l} \right)-\ln\left( \bar{a} \right)}ln(\bar{l})$$

The total energy in normalized form is

$$\bar{U}_{total}=\bar{U}_{adhesion}+\bar{U}_{tension}=-\int_{0}^{\bar{a}} 2\pi\bar{r}\cdot\sqrt{1+\bar{w}_{I}^{'2}}d\bar{r}+\gamma(\int_{0}^{\bar{a}} 2\pi\bar{r}\cdot\sqrt{1+\bar{w}_{I}^{'2}}d\bar{r}-\pi\bar{a}^{2}+\int_{\bar{a}}^{\bar{L}} 2\pi\bar{r}\cdot\sqrt{\left( 1+\bar{w}_{II}^{'2} \right)}d\bar{r}-\pi\left( \bar{L}^{2}-\bar{a}^{2} \right)) (S45)$$

Carrying out the integrals gives:

$$\bar{U}_{total}=-\left( \frac{2}{3}\pi\left( 1+\bar{a}^{2} \right)^{\frac{3}{2}}-\frac{2}{3}\pi\right)+\gamma\left( \frac{2}{3}\pi\left( 1+\bar{a}^{2} \right)^{\frac{3}{2}}-\frac{2}{3}\pi-\pi\bar{a}^{2}+\frac{\pi\sqrt{1+\frac{\bar{c}_{1}^{2}}{\bar{l}^{2}}}\bar{l}\left( \bar{l}\sqrt{\bar{c}_{1}^{2}+\bar{l}^{2}}+\bar{c}_{1}^{2}\ln\left( \bar{l}+\sqrt{\bar{c}_{1}^{2}+\bar{l}^{2}} \right) \right)}{\sqrt{\bar{c}_{1}^{2}+\bar{l}^{2}}}-\frac{\pi\sqrt{1+\frac{\bar{c}_{1}^{2}}{\bar{a}^{2}}}\bar{a}\left( \bar{a}\sqrt{\bar{c}_{1}^{2}+\bar{a}^{2}}+\bar{c}_{1}^{2}\ln\left( \bar{a}+\sqrt{\bar{c}_{1}^{2}+\bar{a}^{2}} \right) \right)}{\sqrt{\bar{c}_{1}^{2}+\bar{a}^{2}}}-\pi\left( \bar{L}^{2}-\bar{a}^{2} \right) \right) (S46)$$

Again, we find $\bar{a}$ by minimization and we calculate the force by

$$\bar{F}= \gamma\frac{d\bar{w}}{d\bar{r}}\cdot2\pi\bar{r}=2\pi\gamma\bar{r}\frac{\bar{c}_{1}}{\bar{r}}=2\pi\gamma\frac{\bar{\delta}+\frac{\bar{a}^{2}}{2}}{\ln\left( \bar{l} \right)-\ln\left( \bar{a} \right)} \left( S47 \right)$$

The results for force-deflection and contact radius as a function of deflection are shown in Fig. 4 (c,d) of the main text. Figure S3 (a) shows how contact width at zero force decreases with tension. As in the 2D case, there is a slow decline and no distinct value at which adhesion is blocked. Fig. S3(b) shows the pull-off force as a function of tension. Again, there is a weak decline in pull off force.

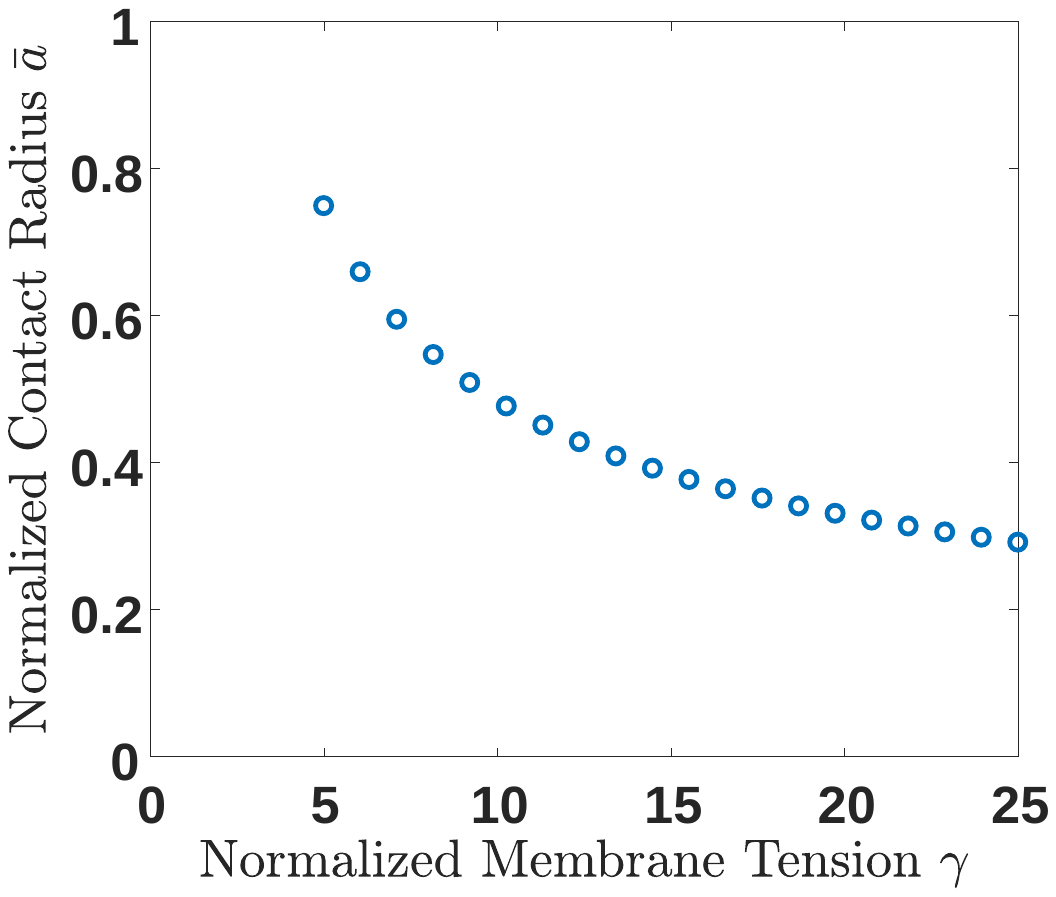

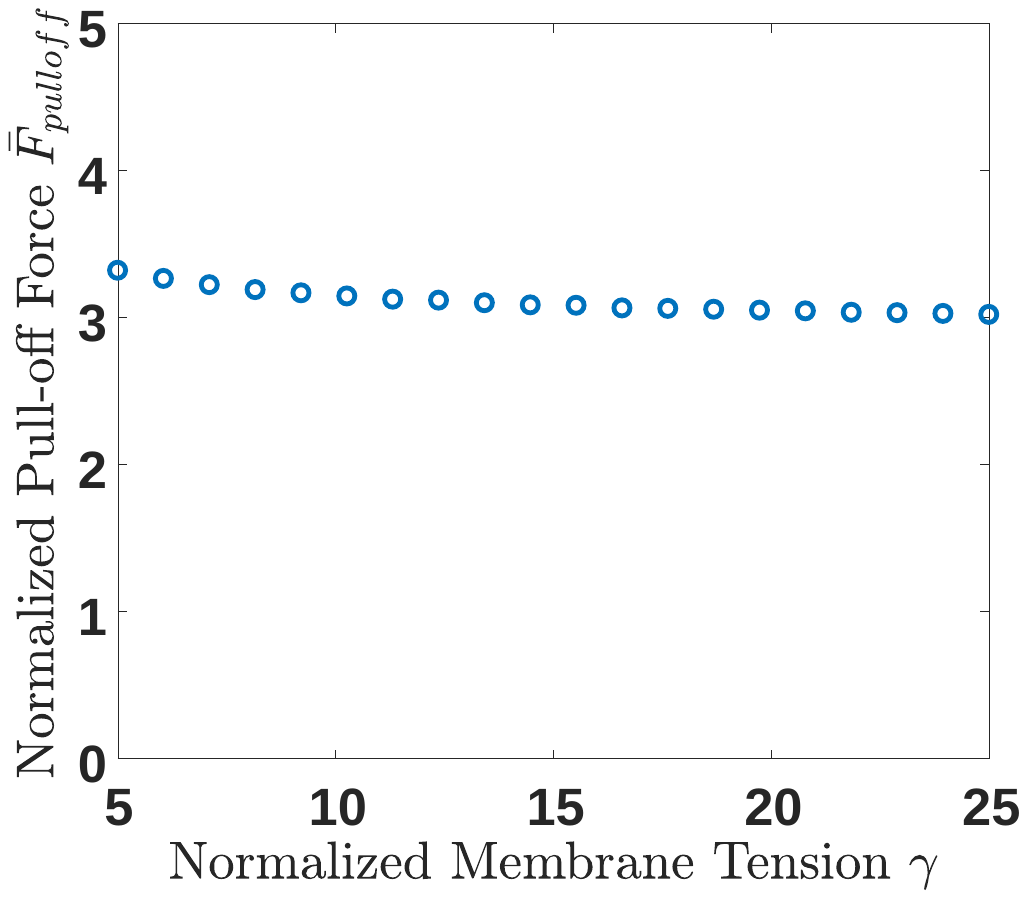

(a)	(b)

Figure S3 (a) Normalized contact width at zero external force as a function of normalized membrane tension. (b) Normalized pull-off force as a function of normalized membrane tension.

**S2.2 Bending limit**

The governing equation is obtained from equation (6) of the main text by setting *γ=0*:

$$\frac{1}{r}\frac{d}{dr}\left\{ r\frac{d}{dr}\left[ \frac{1}{r}\frac{d}{dr}\left( r\frac{dw}{dr} \right) \right] \right\}=0 (S48)$$

The solution in region I remains unchanged. The solution in region II is

$$\bar{w}\left( \bar{r} \right)=\bar{c}_{1}\log\left( \bar{r} \right)+\bar{c}_{2}\cdot\frac{1}{2}\bar{r}^{2}+\bar{c}_{3}\left( \frac{1}{2}\bar{r}^{2}\log\left( \bar{r} \right)-\frac{1}{4}\bar{r}^{2} \right)+\bar{c}_{4} (S49)$$

Applying the boundary and matching conditions

$$\bar{w}_{I}\left( \bar{a} \right)=\bar{\delta}+\frac{\bar{a}^{2}}{2} (S50)$$

$$\bar{w}_{I}'\left( \bar{a} \right)=\bar{a} (S51)$$

$$\bar{w}_{II}\left( \bar{l} \right)=0 (S52)$$

$$\bar{w}_{II}'\left( \bar{l} \right)=0 (S53)$$

we obtain

$\bar{c}_{1}=\frac{\bar{l}^{2}(-\bar{a}^{4}+\bar{a}^{2}\bar{l}^{2}-4\bar{a}^{2}\bar{\delta}\log\left( \bar{a} \right)+4\bar{a}^{2}\bar{\delta}\log\left( \bar{l} \right))}{\bar{a}^{4}-2\bar{a}^{2}\bar{l}^{2}+\bar{l}^{4}-4\bar{a}^{2}\bar{l}^{2}{\log\left( \bar{a} \right)}^{2}+8\bar{a}^{2}\bar{l}^{2}\log\left( \bar{a} \right)\log\left( \bar{l} \right)-4\bar{a}^{2}\bar{l}^{2}{\log\left( \bar{l} \right)}^{2}}$	(S54)

$\bar{c}_{2}=\frac{\bar{a}^{4}-\bar{a}^{2}\bar{l}^{2}+4\bar{a}^{2}\bar{\delta}\log\left( \bar{a} \right)-4\bar{\delta}\bar{l}^{2}\log\left( \bar{l} \right)+4\bar{a}^{2}l^{2}\log\left( \bar{a} \right)\log\left( \bar{l} \right)-4\bar{a}^{2}l^{2}{log(\bar{l})}^{2}}{{-\bar{a}}^{4}+2\bar{a}^{2}\bar{l}^{2}-\bar{l}^{4}+4\bar{a}^{2}\bar{l}^{2}{\log\left( \bar{a} \right)}^{2}-8\bar{a}^{2}\bar{l}^{2}\log\left( \bar{a} \right)\log\left( \bar{l} \right)+4\bar{a}^{2}\bar{l}^{2}{\log\left( \bar{l} \right)}^{2}}$	(S55)

$\bar{c}_{3}=\frac{4(\bar{a}^{2}\bar{\delta}-\bar{\delta}\bar{l}^{2}+\bar{a}^{2}l^{2}\log\left( \bar{a} \right)-\bar{a}^{2}l^{2}log(\bar{l}))}{{-\bar{a}}^{4}+2\bar{a}^{2}\bar{l}^{2}-\bar{l}^{4}+4\bar{a}^{2}\bar{l}^{2}{\log\left( \bar{a} \right)}^{2}-8\bar{a}^{2}\bar{l}^{2}\log\left( \bar{a} \right)\log\left( \bar{l} \right)+4\bar{a}^{2}\bar{l}^{2}{\log\left( \bar{l} \right)}^{2}}$	(S56)

$\bar{c}_{4}=\frac{\bar{l}^{2}(\bar{a}^{4}+2\bar{a}^{2}\bar{\delta}-\bar{a}^{2}\bar{l}^{2}-2\bar{\delta}\bar{l}^{2}+4\bar{a}^{2}\bar{\delta}\log\left( \bar{a} \right)+2\bar{a}^{2}\bar{l}^{2}\log\left( \bar{a} \right)-2\bar{a}^{4}\log\left( \bar{l} \right)-4\bar{a}^{2}\bar{\delta}\log\left( \bar{l} \right)-8\bar{a}^{2}\bar{\delta}\log\left( \bar{a} \right)\log\left( \bar{l} \right)+8\bar{a}^{2}\bar{\delta}{log(\bar{l})}^{2})}{2(\bar{a}^{4}-2\bar{a}^{2}\bar{l}^{2}+\bar{l}^{4}-4\bar{a}^{2}\bar{l}^{2}{\log\left( \bar{a} \right)}^{2}+8\bar{a}^{2}\bar{l}^{2}\log\left( \bar{a} \right)\log\left( \bar{l} \right)-2\bar{a}^{2}\bar{l}^{2}{log(\bar{l})}^{2})}$	(S57)

The total free energy is:

$$\bar{U}_{total}=\bar{U}_{adheison}+\bar{U}_{bending}=-\int_{0}^{\bar{a}} 2\pi\bar{r}\cdot\sqrt{1+\bar{w}_{I}^{'2}}d\bar{r}+ \int_{0}^{\bar{a}} \pi\alpha\left( \frac{\partial^{2}\bar{w}_{I}}{\partial\bar{r}^{2}}+\frac{1}{\bar{r}}\frac{\partial\bar{w}_{I}}{\partial\bar{r}} \right)^{2}\bar{r}d\bar{r}+\int_{\bar{a}}^{\bar{l}} \pi\alpha\left( \frac{\partial^{2}\bar{w}_{II}}{\partial\bar{r}^{2}}+\frac{1}{\bar{r}}\frac{\partial\bar{w}_{II}}{\partial\bar{r}} \right)^{2}\bar{r}d\bar{r} (S58)$$

Or,

$$\bar{U}_{total}=-\frac{2\pi}{3}(\left( 1+\bar{a}^{2} \right)^{\frac{3}{2}}-1)+\pi\alpha(\frac{\bar{a}^{2}}{2}+\frac{2\bar{a}^{3}}{3}+\frac{\bar{a}^{4}}{4})+\frac{\pi\alpha}{12\bar{l}^{2}}\left( 3\left( 8\bar{c}_{1}\bar{c}_{3}\left( -2\bar{c}_{2}+\bar{c}_{3} \right)\bar{l}^{2}+\left( 8{\bar{c}_{2}}^{2}+\bar{c}_{2}\bar{c}_{3}+{\bar{c}_{3}}^{2} \right)\bar{l}^{4}+{\bar{c}_{1}}^{2}\left( -2+8\bar{c}_{3}\bar{l} \right) \right)+6\bar{l}\left( 4{\bar{c}_{1}}^{2}\bar{c}_{3}+\bar{c}_{3}\left( 4\bar{c}_{2}+\bar{c}_{3} \right)\bar{l}^{3}+8\bar{c}_{1}\bar{c}_{2}\bar{l}\left( -1+\bar{c}_{3}\bar{l} \right)-4\bar{c}_{1}\bar{c}_{3}\bar{l}\left( 1+\bar{c}_{3}\bar{l} \right) \right)\log\left( \bar{l} \right)+6\bar{c}_{3}\bar{l}^{2}\left( \bar{c}_{3}\bar{l}^{2}+\bar{c}_{1}\left( -2+4\bar{c}_{3}\bar{l} \right) \right){\log\left( \bar{l} \right)}^{2}+4{\bar{c}_{1}}^{2}{\bar{c}_{3}}^{2}\bar{l}^{2}{\log\left( \bar{l} \right)}^{3} \right)-\frac{\pi\alpha}{12\bar{a}^{2}}\left( 3\left( 8\bar{c}_{1}\bar{c}_{3}\left( -2\bar{c}_{2}+\bar{c}_{3} \right)\bar{a}^{2}+\left( 8{\bar{c}_{2}}^{2}+\bar{c}_{2}\bar{c}_{3}+{\bar{c}_{3}}^{2} \right)\bar{a}^{4}+{\bar{c}_{1}}^{2}\left( -2+8\bar{c}_{3}\bar{a} \right) \right)+6\bar{a}\left( 4{\bar{c}_{1}}^{2}\bar{c}_{3}+\bar{c}_{3}\left( 4\bar{c}_{2}+\bar{c}_{3} \right)\bar{a}^{3}+8\bar{c}_{1}\bar{c}_{2}\bar{a}\left( -1+\bar{c}_{3}\bar{a} \right)-4\bar{c}_{1}\bar{c}_{3}\bar{a}\left( 1+\bar{c}_{3}\bar{a} \right) \right)\log\left( \bar{a} \right)+6\bar{c}_{3}a^{2}\left( \bar{c}_{3}\bar{a}^{2}+\bar{c}_{1}\left( -2+4\bar{c}_{3}\bar{a} \right) \right){\log\left( \bar{a} \right)}^{2}+4{\bar{c}_{1}}^{2}{\bar{c}_{3}}^{2}\bar{a}^{2}{\log\left( \bar{a} \right)}^{3} \right) (S59)$$

Applied force equals

$$\bar{F}=-\alpha\frac{\partial}{\partial\bar{r}}\left( \Delta\bar{w} \right)\cdot2\pi\bar{r}=-2\bar{\alpha c}_{3}\frac{1}{\bar{r}}\cdot2\pi\bar{r}=-4\bar{\alpha}\pi\bar{c}_{3} (S60)$$

The results are shown in Figs 4(a) and (b) of the main text. Figs S4 (a,b) show the normalized contact radius at zero force and the pull-off force as a function of normalized bending stiffness. Like the 2D case, adhesion is blocked at a critical value of one.

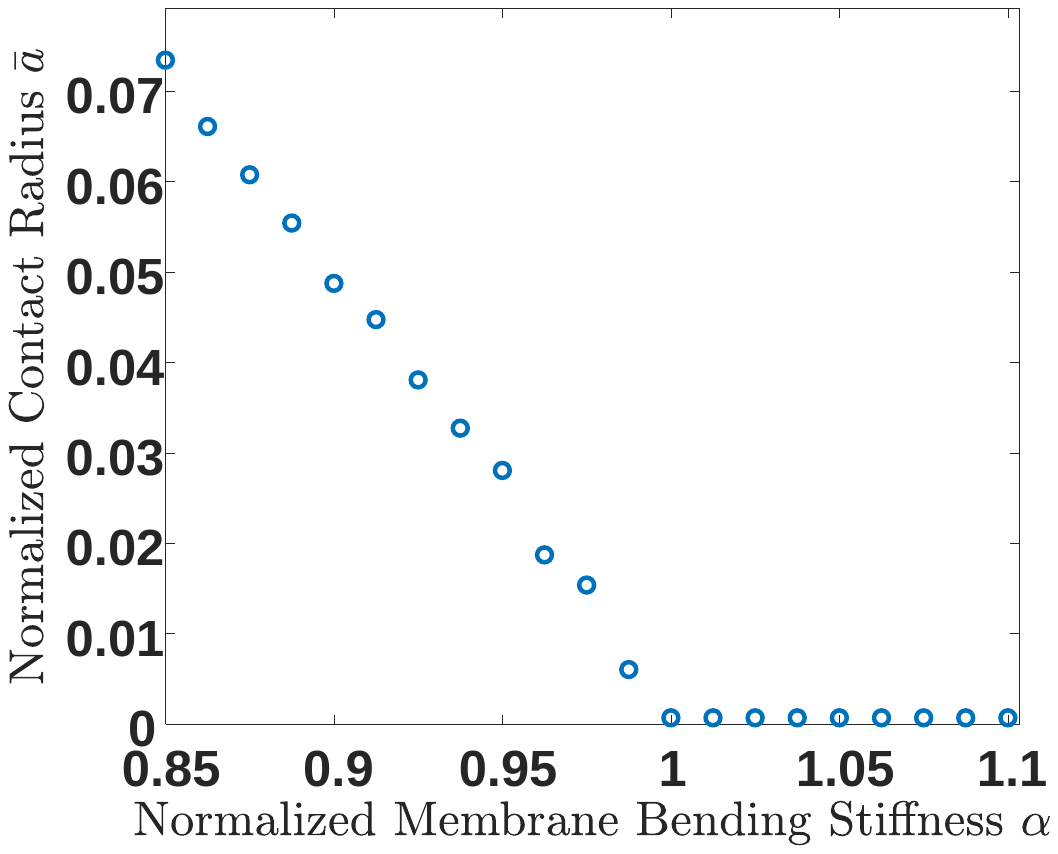

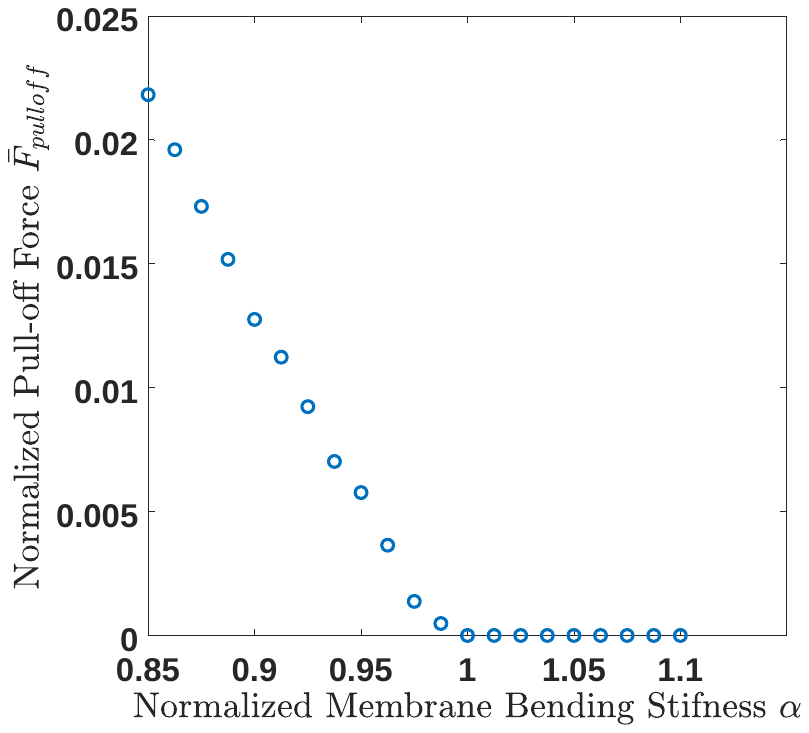

Figure S4 (a) Contact radius at zero external force as a function of normalized membrane bending stiffness. (b) Pull-off force as a function of normalized bending stiffness.

**S2.3 Both bending and tension**

The governing equation for the shape in region II is eq. (6) of the main text. It’s solution is

$$\bar{w}\left( \bar{r} \right)=c_{1}\cdot I_{0}\left( \sqrt{\frac{\gamma\bar{r}^{2}}{\alpha}} \right)+c_{2}\cdot K_{0}\left( \sqrt{\frac{\gamma\bar{r}^{2}}{\alpha}} \right)+c_{3}\cdot\log\left( \bar{r} \right)+c_{4} (S61)$$

in which the unknown constants are determined by applying the same boundary and matching conditions as in section 2.2. We have obtained explicit results but they are extremely lengthy and so won’t be reproduced here. They are available as Mathematica® source files from the authors upon request.

The total free energy on the membrane is calculated by

$$\bar{U}_{total}=\bar{U}_{adhesion}+\bar{U}_{tension}+\bar{U}_{bending}=-\int_{0}^{\bar{a}} 2\pi\bar{r}\cdot\sqrt{1+\bar{w}_{I}^{'2}}d\bar{r}+\gamma\left( \int_{0}^{\bar{a}} 2\pi\bar{r}\cdot\sqrt{1+\bar{w}_{I}^{'2}}d\bar{r}-\pi\bar{a}^{2}+\int_{\bar{a}}^{\bar{L}} 2\pi\bar{r}\cdot\sqrt{\left( 1+\bar{w}_{II}^{'2} \right)}d\bar{r}-\pi\left( \bar{L}^{2}-\bar{a}^{2} \right) \right)+\int_{0}^{\bar{a}} \pi\alpha\left( \frac{\partial^{2}\bar{w}_{I}}{\partial\bar{r}^{2}}+\frac{1}{\bar{r}}\frac{\partial\bar{w}_{I}}{\partial\bar{r}} \right)^{2}\bar{r}d\bar{r}+\int_{\bar{a}}^{\bar{l}} \pi\alpha\left( \frac{\partial^{2}\bar{w}_{II}}{\partial\bar{r}^{2}}+\frac{1}{\bar{r}}\frac{\partial\bar{w}_{II}}{\partial\bar{r}} \right)^{2}\bar{r}d\bar{r} (S62)$$

and the normalized external force is

$$\bar{F}=2\pi\bar{r}\left( -\alpha\frac{\partial}{\partial\bar{r}}\left( \Delta\bar{w} \right)+\gamma\frac{d\bar{w}}{d\bar{r}} \right) (S63)$$
